# The chromatin factor Gon4l regulates embryonic axis extension by promoting mediolateral cell polarity and notochord boundary formation through negative regulation of cell adhesion

**DOI:** 10.1101/154310

**Authors:** Margot L K Williams, Atsushi Sawada, Terin Budine, Chunyue Yin, Paul Gontarz, Lilianna Solnica- Krezel

## Abstract

Anteroposterior axis extension during vertebrate gastrulation requires cell proliferation, embryonic patterning, and morphogenesis to be spatiotemporally coordinated, but the underlying genetic mechanisms remain poorly understood. Here we define a role for the conserved chromatin factor Gon4l, encoded by *ugly duckling (udu),* in coordinating tissue patterning and axis extension during zebrafish gastrulation. Although identified as a recessive enhancer of short axis phenotypes in planar cell polarity (PCP) mutants, we found that Gon4l functions in a genetically independent, partially overlapping fashion with PCP signaling to regulate mediolateral cell polarity underlying axis extension in part by promoting notochord boundary formation. We identified direct genomic targets of Gon4l and found that it acts as both a positive and negative regulator of gene expression, including limiting expression of the cell-cell and cell-matrix adhesion molecules EpCAM and Integrinα3b. Excess *epcam* or *itga3b* in wild-type gastrulae phenocopied notochord boundary defects of *udu* mutants, while downregulation of *itga3b* suppressed them. By promoting formation of this anteroposteriorly aligned boundary and associated cell polarity, Gon4l cooperates with PCP signaling to coordinate morphogenesis with the anteroposterior embryonic axis.

---

Gastrulation is a critical period of animal development during which the three primordial germ layers - ectoderm, mesoderm, and endoderm - are specified, patterned, and shaped into a rudimentary body plan. During vertebrate gastrulation, mesoderm and endoderm become internalized to underlie ectoderm and the three germ layers thin and expand through epibolic movements. The hallmark of the vertebrate body plan is an elongated anteroposterior (AP) axis that emerges as the result of convergence and extension (C&E), a conserved set of gastrulation movements characterized by the concomitant AP elongation and mediolateral (ML) narrowing of the germ layers^1-4^. C&E is accomplished by a combination of polarized cell behaviors, including directed migration and ML intercalation behavior (MIB)^5^^-^^7^. During MIB, cells elongate and align their bodies and protrusions in the ML dimension and intercalate preferentially between their anterior and posterior neighbors^7^. This polarization of cell behaviors with respect to the AP axis is regulated by the planar cell polarity (PCP) and other signaling pathways^8^^-^^12^. Because these pathways are essential for MIB and C&E but do not affect cell fates^12^^-^^14^, other mechanisms must spatiotemporally coordinate morphogenesis with embryonic patterning to ensure normal development. BMP, for example, coordinates dorsal-ventral axis patterning with morphogenetic movements by limiting expression of PCP signaling components and C&E to the embryo’s dorsal side^15^. In general, though, molecular mechanisms that coordinate gastrulation cell behaviors with axial patterning are poorly understood, and remain one of the key outstanding questions in developmental biology.

Epigenetic regulators offer a potential mechanism by which broad networks of embryonic patterning and morphogenesis genes can be co-regulated, conceivably by altering chromatin state within embryonic cells to regulate which subsets of genes are available for transcription. Epigenetic modifiers like Histone acetyl transferases (HATs), deacetylases (HDACs), and methyltransferases associate within protein complexes containing chromatin factors that are thought to regulate their binding at specific genomic regions in context-specific ways^16^. The identities, functions, and specificity of chromatin factors with roles during embryogenesis are only now being elucidated, and often have described roles in cell fate specification and embryonic patterning^17^^-^^19^. The contribution of epigenetic regulation to gastrulation cell movements, however, is particularly poorly understood.

Here, we describe the chromatin factor Gon4l, encoded by *ugly duckling (udu),* as a novel regulator of embryonic axis extension during zebrafish gastrulation. *udu* was identified in a forward genetic screen for enhancers of short axis phenotypes in PCP mutants, but we found it functions in parallel to PCP signaling. Instead, complete maternal and zygotic *udu* (MZ*udu*) deficiency produced a distinct set of morphogenetic and cell polarity phenotypes that implicate the notochord boundary in ML cell polarity and cell intercalation during C&E. Extension defects in MZ*udu* mutants were remarkably specific, as internalization, epiboly and convergence gastrulation movements occurred normally. Gene expression profiling revealed that Gon4l regulates expression of a large portion of the zebrafish genome, including housekeeping genes, patterning genes, and many with known or potential roles in morphogenesis. Furthermore, Gon4l-associated genomic loci were identified by DNA adenine methyltransferase identification^20, 21^ paired with high throughput sequencing (DamID-seq), revealing both positive and negative regulation of putative direct targets by Gon4l. Mechanistically, we found that increased expression of *epcam* and *itga3b*, direct targets of Gon4l-dependent repression during gastrulation, were largely causative of notochord boundary defects in MZ*udu* mutants. This report thereby defines a critical role for a chromatin factor in the regulation of gastrulation cell behaviors in vertebrate embryos: by ensuring proper formation of the AP-aligned notochord boundary and associated ML cell polarity, Gon4l cooperates with PCP signaling to coordinate morphogenesis that extends the AP embryonic axis.

## RESULTS

### Gon4l is a novel regulator of axis extension during zebrafish gastrulation

To identify novel regulators of C&E, we performed a three generation synthetic mutant screen^22, 23^ using zebrafish carrying a hypomorphic allele of the planar cell polarity (PCP) gene *knypek (kny)/glypican 4, kny^m818^*^10^. F0 wild-type (WT) males were mutagenized with *N*-ethyl, *N*-nitrosourea (ENU) and outcrossed to WT females. The resulting F1 fish were outcrossed to *kny^m818/818^* males (rescued by *kny* RNA injection) to generate F2 families whose F3 offspring were screened during early segmentation (11-14 hours post fertilization (hpf)) and late embryogenesis (24hpf) for short axis phenotypes (Fig.1a). Screening nearly 100 F2 families yielded eight recessive mutations that enhanced axis elongation defects in *kny^m818/818^* embryos. *vu68* was found to be a new L227P *kny* allele, and *vu64* was a new Y219* nonsense allele of the core PCP gene *trilobite(tri)/vangl2*, mutations in which exacerbate *kny* mutant phenotypes^24^, demonstrating effectiveness of our screening strategy. We focused on *vu66/vu66* mutants, which displayed a pleiotropic phenotype at 24hpf, including shortened AP axes, reduced tail fins, and heart edema (Fig.1d, Supplemental Fig.1b).

**Figure 1:**
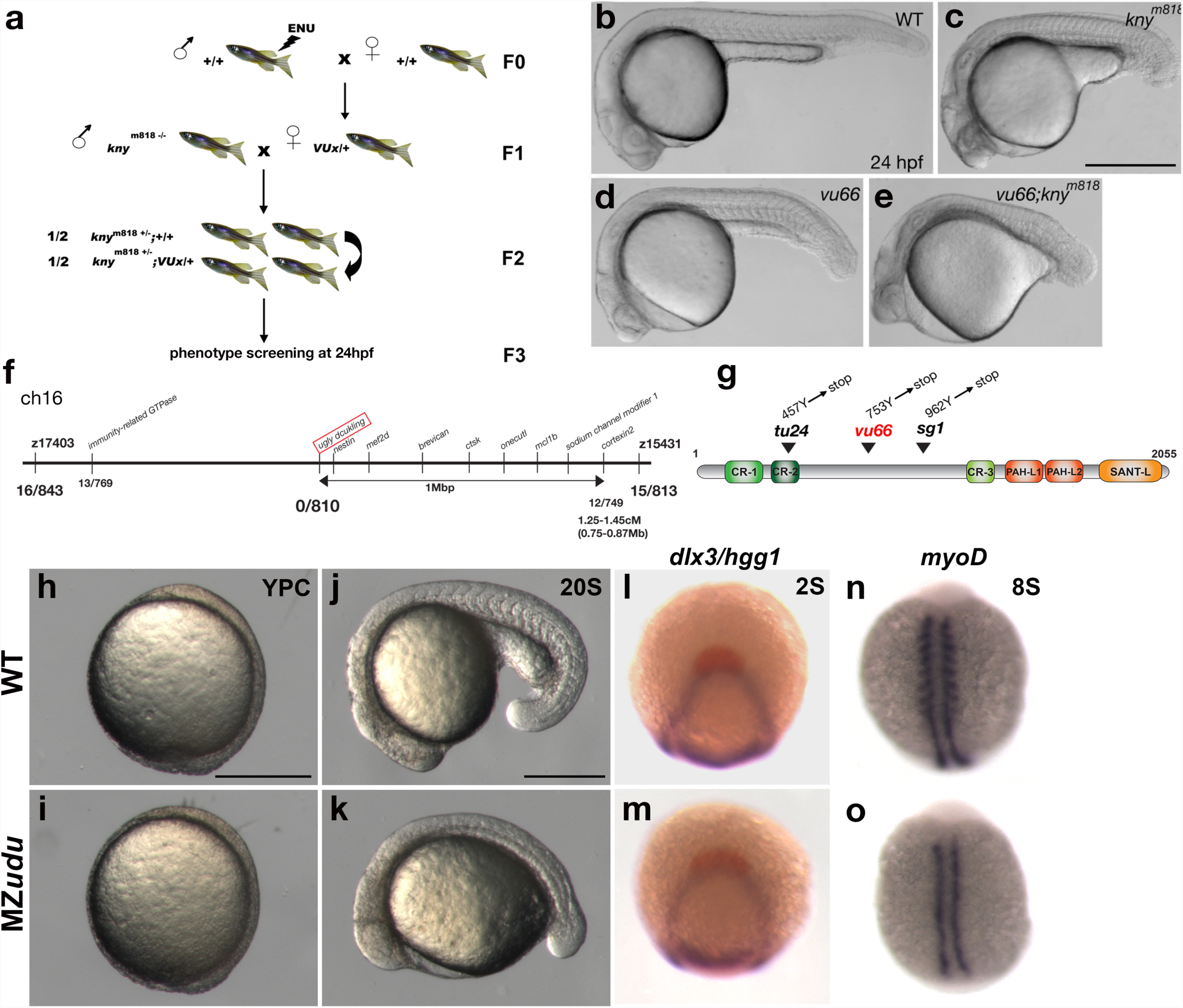
A forward genetic screen identifies *ugly duckling (udu)/gon4l* as a novel regulator of axis extension in zebrafish embryos. **a)** Schematic of a synthetic screen to identify enhancers of the short axis phenotype in *kny^m818/m818^* zebrafish mutants. **b-e)** Phenotypes at 24 hpf: wild-type (WT) **(b)**, *kny^m818/m818^* (c), *vu66/vu66* (d), *kny^m818/m818^; vu66/vu66* compound mutants (e). **f)** Diagram of mapping *vu66* mutation to Chromosome 16. Bold numbers below specify the number of recombination events between *vu66* and the indicated loci. **g)** Diagram of the Gon4l protein encoded by the *ugly duckling (udu)* locus. Arrowheads indicate residues mutated in *vu66* and other described *udu* alleles. **h-k)** Live WT (h,j) and maternal zygotic (MZ)udu (i,k) embryos at yolk plug closure (YPC) (h,i) and 20 somite stage (j,k). **l-m**) Whole mount *in situ* hybridization (WISH) for *dlx3* (purple) and *hgg1* (red) in WT (l) and MZ*udu* (m) embryos at 2 somite stage. **n-o**) WISH for *myoD* in WT (n) and MZ*udu* (o) embryos at 8 somite stage. Anterior is to the left in b-e, j-k; anterior is up in h-i, l-o. Scale bar is 500 μm in b-e and 300 μm in h-o.

Employing simple sequence repeat mapping strategy^25,26^, we mapped the *vu66* mutation to a small region on Chromosome 16 that contains the *ugly duckling* (*udu*) gene (Fig.1b). *udu* encodes a conserved chromatin factor homologous to *gon4* in *C. elegans* and the closely related *Gon4l* in mammals^27^. The previously described *udu* mutant phenotypes resemble those of homozygous *vu66* embryos, including a shorter body axis and fewer blood cells^27, 28^, making *udu* an excellent candidate for this novel PCP enhancer. Sequencing cDNA of the *udu* coding region from 24hpf *vu66/vu66* embryos revealed a T to A transversion at position 2,261 predicted to change 753Y to a premature STOP codon (Fig.1g). Furthermore, *vu66* failed to complement a known *udu^sq^*^1^ allele^27^ (Supplemental Fig.1c). Together, these data establish *vu66* as a new *udu* allele and identify it as a recessive enhancer of axis extension defects in *kny* PCP mutant gastrulae.

### Complete loss of maternal and zygotic *udu* function impairs axial extension

A majority of zebrafish genes are expressed maternally^29, 30^, and their expression can mask the effect of zygotic mutations on embryo patterning and gastrulation^31, 32^. Because *udu* is highly maternally expressed^27^, we employed germline replacement^33^ to generate otherwise WT females carrying *udu^vu66/vu66^* (*udu-/-*) germline, and therefore lacking maternal *udu* expression. Females harboring *udu^vu66/vu66^*germline were crossed to *udu^vu66/+^* or *udu^vu66/vu66^* germline males to produce 50% or 100% embryos lacking both maternal (M) and zygotic (Z) *udu* function, respectively, hereafter referred to as MZ*udu* mutants. Strict maternal loss of *udu* produced no obvious phenotypes (Fig.4a). *MZudu-/-* embryos appeared outwardly to develop normally until mid-gastrula stages (Fig.1i), specified the three germ layers (Fig.1m, Supplemental Fig.2), formed an embryonic shield marked by *gsc* expression (Supplemental Fig.2c,d), and completed epiboly on schedule (Fig.1i). However, they exhibited clear abnormalities at the onset of segmentation, as somites were largely absent in mutants (Fig.1k,o). Although *myoD* expression was observed within adaxial cells by whole mount *in situ* hybridization (WISH), its expression was not detected in nascent somites (Fig. 1o). Formation of adaxial cells is consistent with normal expression of their inducer *shh* in the axial mesoderm^34^ (Supplemental Fig.2h), and *ntla/brachyury* expression in the axial mesoderm was also largely intact (Supplemental Fig.2j). Importantly, *MZudu-/-* embryos were markedly shorter than age-matched WT controls throughout segmentation (Fig.1k) and at 24hpf (Supplemental Fig. 3e,h, Fig.4b). Although increased cell death was observed in MZ*udu* and *Zudu* mutants^35^, inhibiting apoptosis within MZ*udu* embryos via injection of RNA encoding the anti-apoptotic mitochondrial protein Bcl-xL^36^ did not suppress their short axis phenotype (Supplemental Fig.3a-f). MZ*udu* mutants displayed a number of other phenotypes, including blood^27, 28^ and cardiac deficiencies, which will be described further elsewhere (Budine, Williams, LSK, unpublished). These results demonstrate a role for Gon4l in axial extension during gastrulation.

**Figure 2:**
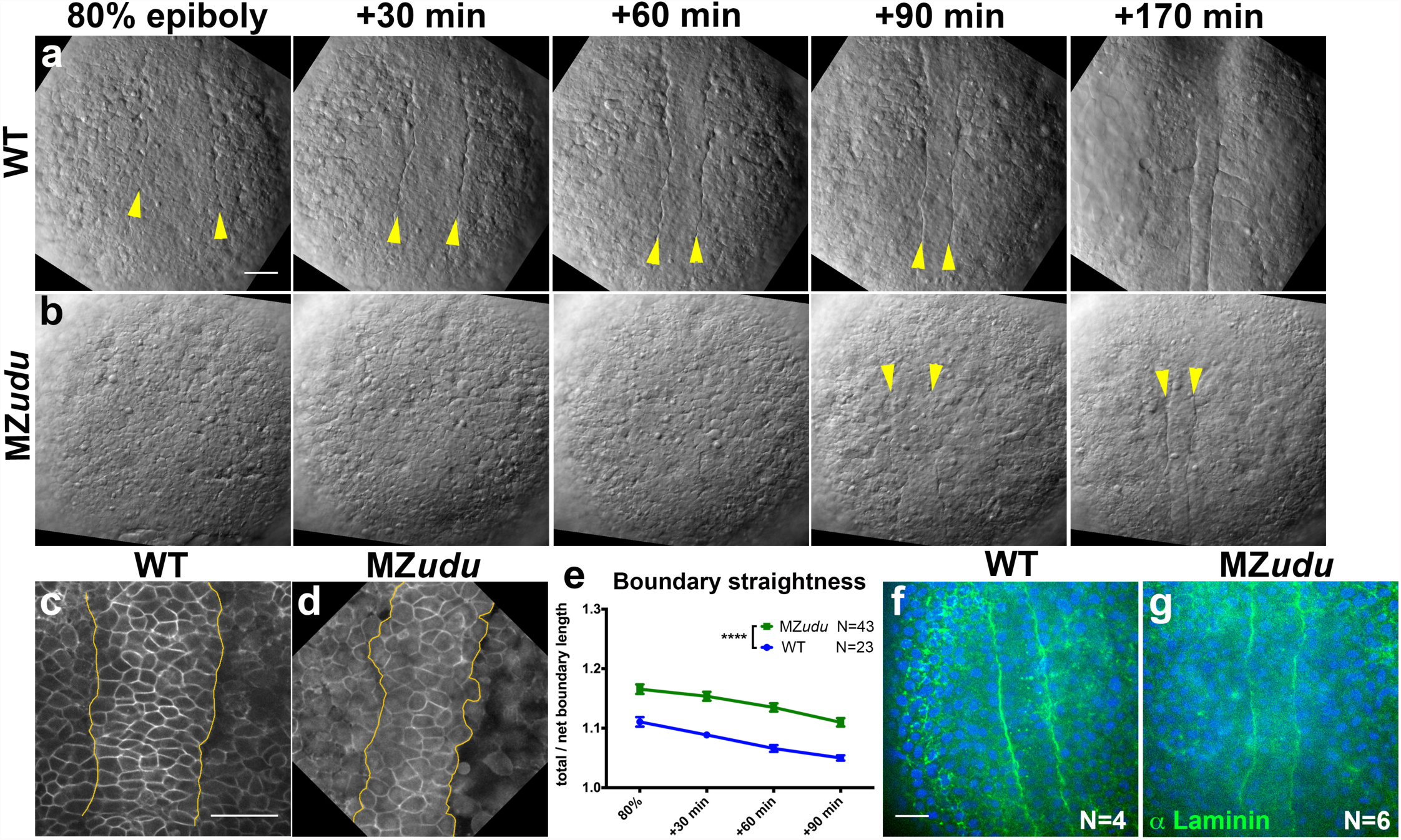
MZ*udu* mutant gastrulae exhibit irregular notochord boundaries. **a-b**) Still images from live Nomarski time-lapse series of the dorsal mesoderm in WT (a) and MZ*udu* embryos (b) at the time points indicated. Arrowheads indicate notochord boundaries. **c-d**) Live confocal microscope images of representative WT (c) and MZ*udu* (d) embryos expressing membrane Cherry. Yellow lines demarcate notochord boundaries. **e**) Quantification of notochord boundary straightness in live WT and MZ*udu*- */-* gastrulae throughout gastrulation. Symbols are means with SEM (2-way ANOVA, ^****^p<0.0001). **f-g**) Confocal microscope image of immunofluorescent staining for pan-Laminin in WT (f) and MZ*udu* (g) embryos at 2 somite stage. Scale bars are 50 μm. Anterior is up in all images.

### Gon4l regulates formation of the notochord boundary

Time-lapse Nomarski (Fig.2a-b) and confocal (Fig.2c-d) microscopy of dorsal mesoderm in *MZudu-/-* gastrulae revealed reduced definition and regularity of the boundary between axial and paraxial mesoderm, hereafter referred to as the notochord boundary, compared to WT (Fig.2a-b, arrowheads). The ratio of the total/net length of notochord boundaries was significantly higher in *MZudu-/-* than in WT gastrulae at all time points (Fig.2e), indicative of decreased straightness. Interestingly, Laminin was detected by immunostaining at the notochord boundary of both WT and *MZudu-/-* embryos (Fig.2f-g), indicating that MZ*udu* mutants form a *bona fide* boundary, albeit an irregular one. These results demonstrate that Gon4l is necessary for proper formation of the notochord boundary during gastrulation.

### Mediolateral polarity and intercalation of axial mesoderm cells are reduced in MZ*udu-/-* gastrulae

In vertebrate gastrulae, C&E is achieved chiefly through ML intercalation of polarized cells that elongate and align their cell bodies with the ML embryonic axis^5, 7, 9, 11^. To determine whether cell polarity defects underlie reduced axis extension in Mz*udu*-/- gastrulae, we measured cell orientation (the angle of a cell’s long axis with respect to the embryo’s ML axis; Fig.3b,e) and cell body elongation (length-to-width or aspect ratio (AR); Fig.3c,f) in confocal time-lapse series of MZ*udu-/-* and WT gastrulae expressing membrane Cherry fluorescent protein (mCherry). Given the irregular notochord boundaries observed in MZ*udu-/-* gastrulae (Fig.2d-e), we examined the time course of cell polarization according to a cell’s position with respect to the boundary: i.e. boundary-adjacent “edge” cells were analyzed separately from those one or two cell diameters away (hereafter −1 and −2, respectively), and so on (See Fig.3). Most WT axial mesoderm cells at 80% epiboly were largely ML oriented and somewhat elongated, but boundary-adjacent “edge” cells were less well oriented (median angle= 24.6°) than cells within any of the internal rows (median angles= 15.7°, 16.6°, 21.1°; Fig.3a-c). However, at the end of the gastrula period 90 minutes later, edge cells became highly aligned and elongated (median angle=13.6°, mean AR=2.2) similar to internal cell rows (median angles= 12.1°, 18.4°; Fig.3a-c). All MZ*udu*-/- axial mesoderm cells exhibited severely reduced ML alignment and elongation at 80% epiboly (Fig.3d-f), but 90 minutes later only the edge cells remained less aligned than WT (median angle=17.0°, mean AR=1.9) (Fig.3e), although aspect ratio of the edge and −1 cells remained reduced (Fig. 3f). These results indicate significantly reduced ML orientation of axial mesoderm cells in MZ*udu-/-* gastrulae, a defect that persisted only in boundary-adjacent cells at late gastrulation. Importantly, this reduction in ML cell polarity was accompanied by significantly fewer cell intercalation events within the axial mesoderm of MZ*udu* mutants compared to WT (Fig.3g-i). As ML intercalation is the key cellular behavior required for vertebrate C&E^5, 7^, we conclude that this is likely the primary cause of axial mesoderm extension defects in MZ*udu* mutants. We also observed significantly fewer mitoses in MZ*udu-/-* gastrulae (Fig.3j-l). Because decreased cell proliferation and the resulting reduced number of axial mesoderm cells has been demonstrated to impair extension in zebrafish^37^, this could also be a contributing factor. Together, reduction of ML cell polarity, ML cell intercalations, and cell proliferation provide a suite of mechanisms resulting in impaired axial extension in MZ*udu-/-* gastrulae. Combined with irregular notochord boundaries observed in MZ*udu* mutants, we hypothesize that this boundary provides a ML orientation cue that contributes to ML polarization of axial mesoderm cells, and that this cue is absent or reduced in MZ*udu-/-* gastrulae.

**Figure 3:**
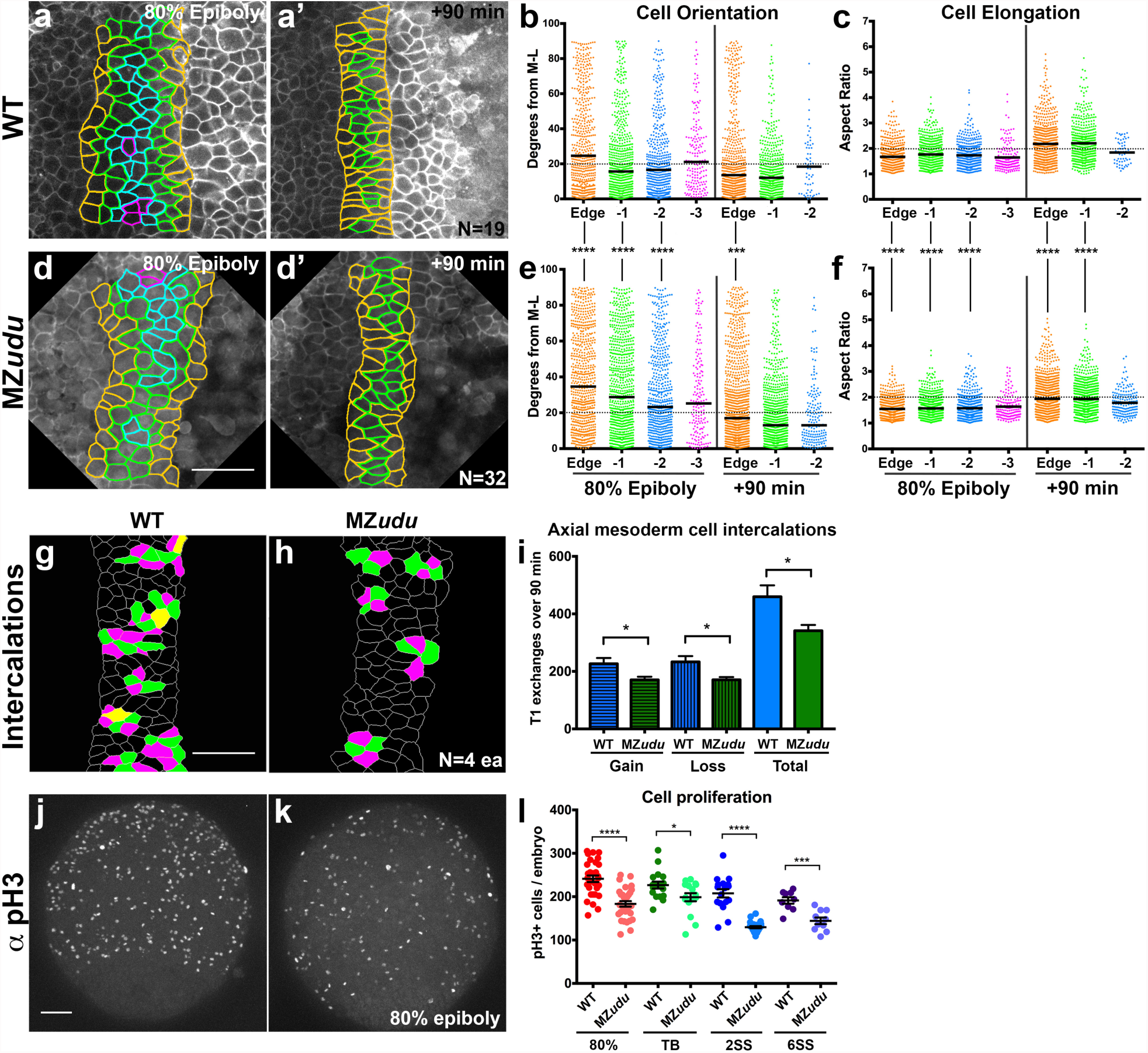
Mediolateral cell polarity and cell intercalations are reduced in the axial mesoderm of MZ*udu-/-* embryos. **a-a’, d-d’**) Still images from live time-lapse confocal movies of the axial mesoderm in WT (a) and MZ*udu-/-* (d) gastrulae at the time points indicated. Cell outlines are colored according to a cell’s position with respect to the notochord boundary. **b, e**) Quantification of axial mesoderm cell orientation at 80% epiboly (left side) and +90 minute (right side) time points. Each dot represents the orientation of the major axis of a single cell with respect to the embryonic ML axis and is colored according to that cell’s position with respect to the notochord boundary (colors match those of the images to the left). Horizontal black lines indicate median values. Asterisks indicate significant differences between WT and MZ*udu* (Kolmogorov-Smirnov test, ^***^p<0.001, ^****^p<0.0001). **c, f**) Quantification of axial mesoderm cell elongation at 80% epiboly (left side) and +90 minute (right side) time points. Each dot represents the aspect ratio of a single cell and is color-coded as in (b). Horizontal black lines indicate mean values. Asterisks indicate significant differences between WT and MZ*udu* (Mann-Whitney test, ^****^p<0.0001). **g-h**) Cell intercalations detected in the axial mesoderm of WT (g) and MZ*udu*-/- gastrulae (h). Cells gaining contacts with neighbors are green, cells losing contacts are magenta, and cells that both gain and lose contacts are yellow. **i**) Quantification of cell intercalation events (T1 exchanges) in WT (blue bars) and MZ*udu-/-* gastrulae (green bars) over 90 minutes. Bars are means with SEM (T test, ^*^p<0.05). **j-k**) 200μm confocal Z projections of immunofluorescent staining for phosphorylated Histone H3 (pH3) in WT (j) and MZ*udu-/-* gastrulae (k) at 80% epiboly. **l**) Quantification of pH3+ cells/embryo at the stages indicated. Each dot represents a single embryo, dark lines are means with SEM (T test, ^*^p<0.05, ^***^p<0.001, ^****^p<0.0001). Scale bars are 50 μm. N indicates the number of embryos analyzed. Anterior/animal pole is up in all images.

### Gon4l regulates axial mesoderm cell polarity and extension independent of PCP signaling

Molecular control of ML cell polarity underlying C&E movements in vertebrate embryos is largely attributed to planar cell polarity (PCP) and Gα12/13 signaling^8^^-^^12^. While zygotic loss of *udu* enhanced axial extension defects in *kny^m818/m818^* PCP mutants (Fig. 1) and axial mesoderm cells in MZ*udu-/-* gastrulae exhibited impaired ML cell polarity and intercalation (Fig.3), it was unclear whether *udu* functions within or parallel to the PCP network. To address this, we generated compound MZ*udu;Zknÿ^fr6/fr6^*(a nonsense/null *kny* allele^10^) mutants utilizing germline replacement as described above. Strikingly, these compound mutant embryos were substantially shorter than single MZ*udu* or *kny^fr6/fr6^* mutants (Fig.4d). Likewise, interference with *vangl2/tri* function in MZ*udu* mutants by injection of MO1-vangl2 antisense morpholino oligonucleotide^38^ also exacerbated axis extension defect of MZ*udu* mutants (Supplemental Fig.4a-d). That reduced levels of PCP components *kny* or *tri* enhanced short axis phenotype resulting from complete *udu* deficiency provides evidence that Gon4l affects axial extension via a parallel pathway. Furthermore, expression domains of genes encoding Wnt/PCP signaling components *kny, tri, wnt5,* and *wnt11* in MZ*udu-/-* gastrulae were comparable to WT (Supplemental Fig.4e-l). Finally, we found that the asymmetric intracellular localization of Prickle (Pk)-GFP, a core PCP component, to anterior cell membranes in WT gastrulae during C&E movements^32, 39^ was not affected in MZ*udu-/-* gastrulae (Fig.4e-f). This is consistent with intact PCP signaling in MZ*udu* mutant gastrulae, and provides further evidence that Gon4l functions largely in parallel to PCP to regulate ML cell polarity and axis extension during gastrulation.

**Figure 4:**
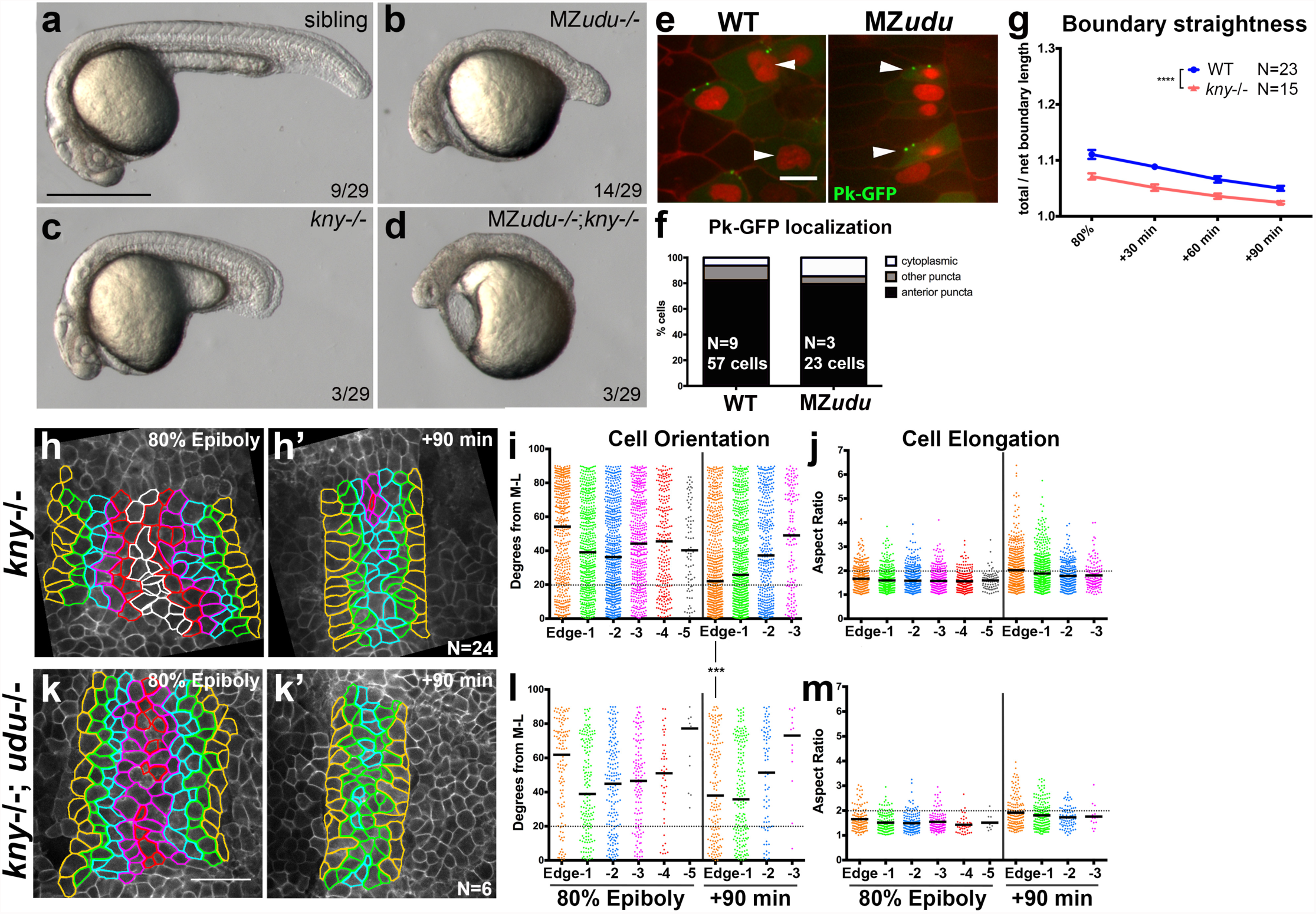
Gon4l regulates axis extension independent of PCP signaling. **a-d**) Live embryos at 24hpf resulting from a cross between a germline-replaced *udu-/-;kny^fr6/+^* female and an *udu+/-;kny^fr6/+^* male. Genotypes are indicated in the upper right corner, the fraction of embryos in the clutch with the pictured phenotype is provided at the bottom. **e**) Mosaically expressed Prickle(Pk)-GFP in WT and MZ*udu*- */-* gastrulae. Arrowheads indicate anteriorly localized Pk-GFP puncta. Membrane Cherry demarcates cell membranes, nuclear-RFP marks cells injected with *pk-gfp* RNA. **f**) Quantification of Pk-GFP localization shown in e. Distribution is not significantly different (chi-square, p=0.07). **g**) Quantification of notochord boundary straightness in WT and *kny-/-* gastrulae. Symbols are means with SEM (2-way ANOVA, ^****^p<0.0001). **h,h’,k,k’**) Still images from live time-lapse confocal movies of the axial mesoderm in *kny-/-* (h) and *kny-/-;udu-/-* (k) gastrulae at the time points indicated. Cell outlines are colored as in Figure 3. **i,l**) Quantification of axial mesoderm cell orientation as in Figure 3 (Kolmogorov-Smirnov test, ^***^p=0.0003). **j,m**) Quantification of axial mesoderm cell elongation as in Figure 3. Scale bar is 500 μm in a-d, 10 μm in e, 50 μm in h-k.

### A Gon4l-dependent boundary cue and PCP signaling cooperate to polarize axial mesoderm cells

In addition to PCP signaling, notochord boundaries are required for proper C&E of the axial mesoderm in ascidian embryos^40^ and are involved in the polarization of intercalating cells during *Xenopus* gastrulation^1^. Boundary defects observed in MZ*udu*-/- gastrulae are not a common feature of mutants with reduced C&E, however, as notochord boundaries in *kny^fr6/fr6^* gastrulae were straighter than in WT siblings (Fig.4g). Consistent with cell polarity defects previously reported in *kny* mutants^10^, all axial mesoderm cells failed to align ML within *kny-/-* embryos at 80% epiboly regardless their position relative to the notochord boundary (Fig.4h-j). After 90 minutes, however, *kny-/-* cells in the edge (and to a lesser extent, −1) position attained distinct ML orientation (median angle= 22.1°). Indeed, the nearer a *kny-/-* cell was to the notochord boundary, the more ML aligned it was likely to be (Fig.4i). This suggests that the notochord boundary provides a ML orientation cue that is independent of PCP signaling and functions across approximately two cell diameters. Furthermore, this boundary-associated cue appears to operate later in gastrulation, whereas PCP-dependent cell polarization is evident by 80% epiboly. Importantly, distinct cell polarity phenotypes observed in MZ*udu* and *kny-/-* gastrulae provide further evidence that Gon4l functions in parallel to PCP signaling.

To assess how PCP signaling interacts with the proposed Gon4l-dependent boundary cue, we examined the polarity of axial mesoderm cells in compound zygotic *kny-/-;udu-/-* mutant gastrulae (Fig.4k-m). As observed in single *kny-/-* mutant gastrulae^10^ (Fig.4h-j), both ML orientation and elongation of all axial mesoderm cells were disrupted in double mutants at 80% epiboly, regardless of position with respect to the notochord boundary. By late gastrulation however, edge cells in double mutant gastrulae failed to attain ML alignment as observed in *kny-/-* mutants, but instead remained largely randomly oriented (median angle= 38.0°) (Fig.4l). This exacerbation of cell orientation defects was correlated with stronger axis extension defects in compound *kny-/-;udu-/-* compared to single *kny-/-* mutants (Fig. 1e). This supports our hypothesis that a Gon4l-dependent boundary cue regulates ML alignment of axial mesoderm cells independent of PCP signaling, and that these two mechanisms function in partially overlapping spatial and temporal domains to cooperatively polarize all axial mesoderm cells and promote axial extension.

### Loss of Gon4l results in large-scale gene expression changes in zebrafish gastrulae

As a nuclear-localized chromatin factor^27, 41^ (Supplemental Fig.3j), Gon4l is unlikely to influence morphogenesis directly. To identify genes regulated by Gon4l with potential roles in morphogenesis, we performed RNA sequencing (RNA-seq) in MZ*udu-/-* and WT tailbud-stage embryos. Analysis of relative transcript levels revealed that more than 11*%* of the genome was differentially expressed (p_adj_<0.01, ≥ 2-fold change) in MZ*udu* mutants (Fig. 5a). Of these ~2,950 differentially expressed genes, 1,692 exhibited increased expression in MZ*udu* compared to 1,259 with decreased expression (Supplemental Table 1). Functional annotation analysis revealed that genes downregulated in MZ*udu*-/- gastrulae were enriched for ontology terms related to chromatin structure, transcription, and translation (Fig.5e), while upregulated genes were enriched for terms related to biosynthesis, metabolism, and protein modifications (Fig.5f). Notably, many of these misregulated genes are considered to have “house-keeping” functions, in that they are essential for cell survival. We further examined genes with plausible roles in morphogenesis, including those encoding signaling and adhesion molecules, and found the majority of genes in both classes were upregulated in MZ*udu-/-* gastrulae (Fig.5c-d). This putative increase in adhesion was of particular interest given the tissue boundary defects observed in MZ*udu* mutants. These findings reveal that although *udu* is ubiquitously expressed in zebrafish embryos^27^, its role in regulating gene expression during gastrulation exhibits some specificity for different classes of encoded proteins, many with known or expected roles in morphogenesis during gastrulation.

**Figure 5:**
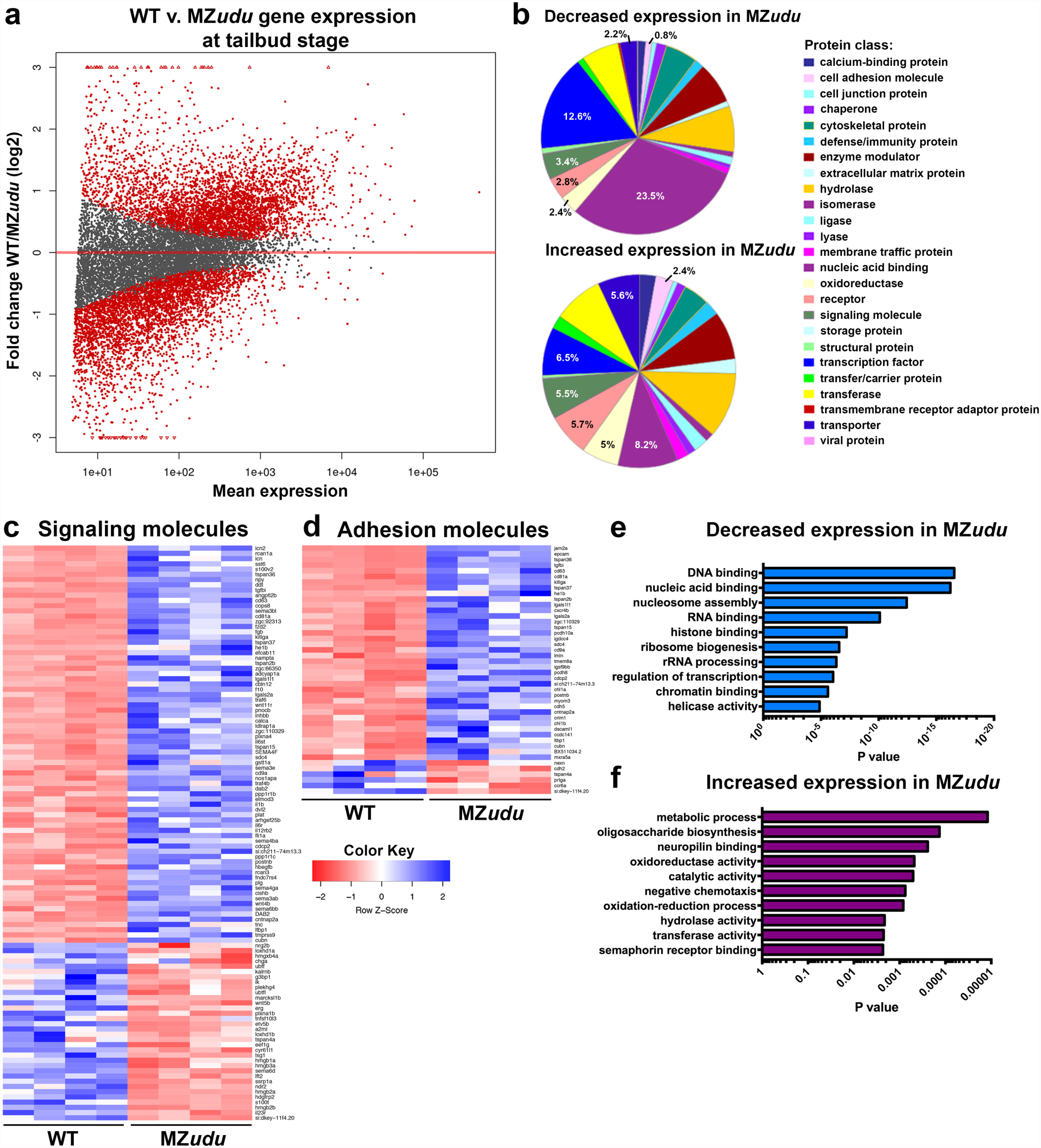
Loss of Gon4l results in large-scale gene expression changes. **a**) Plot of gene expression changes in MZ*udu* mutants compared to WT at tailbud stage as assessed by RNA-sequencing. Red dots indicate genes expressed at significantly different levels than WT (p≤0.01, at least 2-fold change in transcript level). **b**) Protein classes encoded by differentially expressed genes with decreased (top graph) or increased (bottom graph) expression in MZ*udu* mutants. Percentages indicate the number of genes within a given class/total number of genes with increased or decreased expression, respectively. **c-d**) Heat maps of differentially expressed genes annotated as encoding signaling (c) or adhesion molecules (d). The four columns represent two biological and two technical replicates for each WT and MZ*udu*. **e-f**) P-values of the top ten most enriched gene ontology (GO) terms among genes with decreased (e) or increased (f) expression in MZ*udu*-/- embryos compared to WT.

### DamID-seq identifies putative direct targets of Gon4l during gastrulation

To determine genomic loci with which Gon4l protein associates, and thereby distinguish direct from indirect targets of Gon4l regulation, we employed DNA adenine methyltransferase (Dam) identification paired with next generation sequencing (DamID-seq)^20, 21^. To this end, we generated Gon4l fused to *E. coli* Dam, which methylates adenine residues within genomic regions in its close proximity^20^. Small equimolar amounts of RNA encoding a Myc-tagged Gon4l-Dam fusion or a Myc-tagged GFP-Dam control were injected into one-celled embryos, and genomic DNA was collected at tailbud stage (see Material and Methods). In support of this Gon4l-Dam fusion being functional, it localized to the nuclei of zebrafish embryos and partially rescued MZ*udu* embryonic phenotypes (Supplemental Fig.5). Methylated, and therefore Gon4l-proximal, genomic regions were then selected using methylation specific restriction enzymes and adaptors, amplified to produce libraries, and subjected to next generation sequencing. Because Dam is highly active, even an untethered version methylates DNA within open chromatin regions^20^, and so libraries generated from embryos expressing GFP-Dam served as controls.

Unbiased genome-wide analysis of DamID reads revealed a significant enrichment of Gon4l-Dam over GFP-Dam in promoter regions, and a significant underrepresentation of Gon4l within intergenic regions (Fig.6a). Although no global difference was detected within gene bodies (Fig.6a), examination of individual loci revealed approximately 4,500 genes and over 2,300 promoters in which at least one region was highly Gon4l-enriched (P_adj_ ≤0.01, ≥4-fold enrichment over GFP controls) (Fig.6c-g, Supplemental Table 1). Of these, approximately 1,000 genes were co-enriched for Gon4l in both the promoter and gene body (Fig.6c,g). Levels of Gon4l enrichment across a gene and its promoter were significantly correlated (Supplemental Fig.5f), indicating co-enrichment or co-depletion for Gon4l at both gene features of many loci. Within gene bodies, we found robust enrichment specifically within 5’ untranslated regions (UTRs) (Fig.6b), consistent with association of Gon4l at or near transcription start sites (Fig.6d). Of the ~2,950 genes differentially expressed in MZ*udu* mutants, approximately 28% (812) were also enriched for Gon4l at the gene body (492), promoter (170), or both (150) (Fig.6c-g), and will hereafter be described as putative direct Gon4l targets. *histh1,* for example, was among the most downregulated genes by RNA-seq in MZ*udu* mutants and was highly enriched for Gon4l at both its promoter and gene body (Fig.6c). By contrast, *tbx6* expression was also reduced in MZ*udu* mutants, but exhibited no enrichment of Gon4l over GFP controls (Fig.6d). Approximately 35% and 53% of genes enriched for Gon4l in only the gene body or only the promoter, respectively, were positively regulated (i.e. downregulated in MZ*udu* mutants), as were 50% of genes co-enriched at both features. Furthermore, among these positively regulated genes, differential expression levels (the degree to which WT expression exceeded MZ*udu*) correlated positively and significantly with Gon4l enrichment levels across both gene bodies and promoters (Fig.6h). A similar correlation was not observed for negatively regulated genes, hence, the highest levels of Gon4l enrichment were associated with positive regulation of gene expression. These results implicate Gon4l as both a positive and negative regulator of gene expression during zebrafish gastrulation.

**Figure 6:**
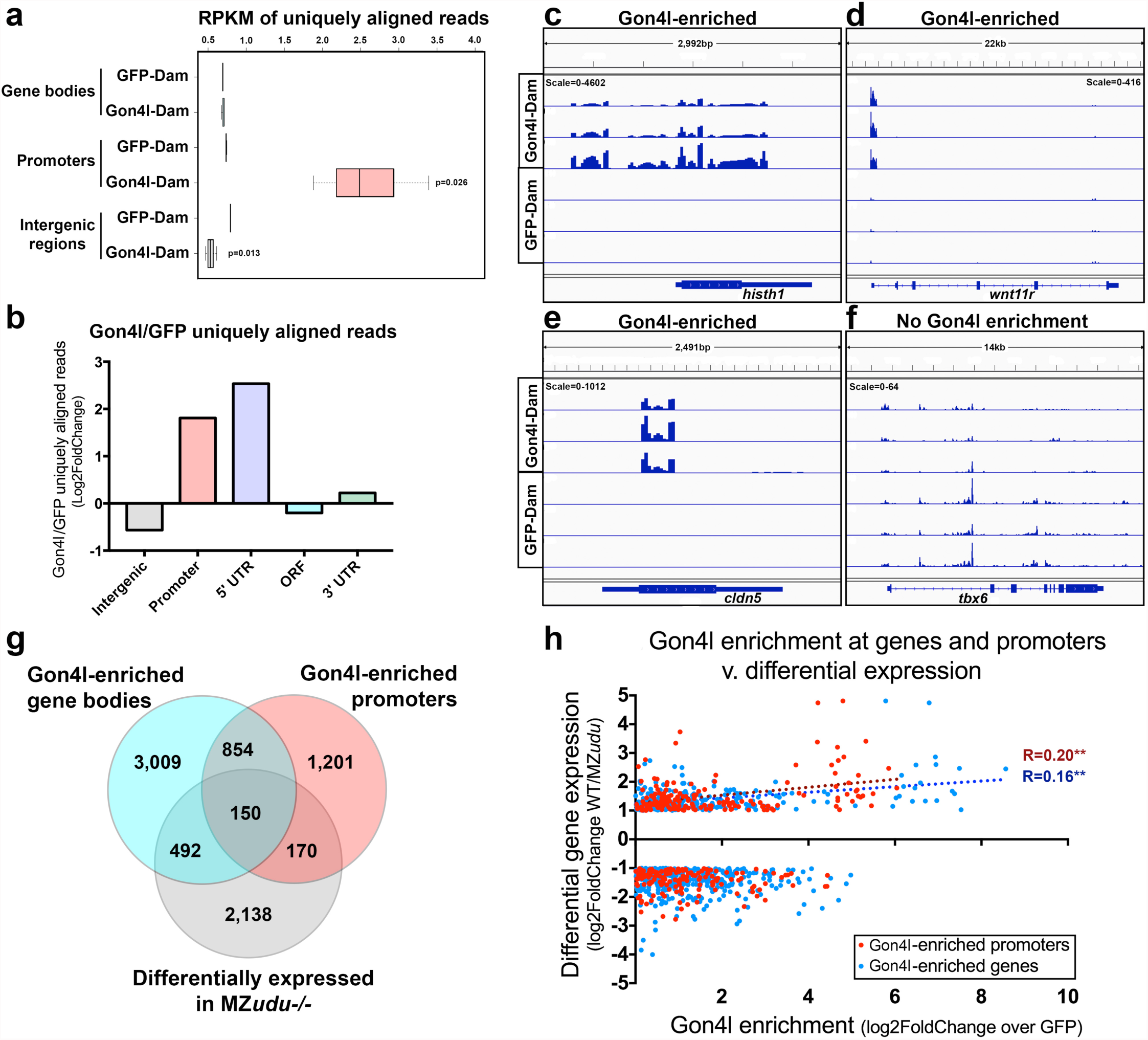
DamID-seq identifies putative direct targets of Gon4l during gastrulation. **a)** Box plot of normalized uniquely aligned DamID reads at each of three categories of genomic regions. P values indicate significant differences between Gon4l-Dam samples and GFP-controls (T-test). **b**) Fold change (log2) Gon4l-Dam over GFP-Dam reads (RPKM) at each of five gene features. **c-f**) Genome browser tracks of Gon4l-Dam and GFP-Dam association at the *histh1* (c), *wnt11r* (d), *cldn5* (e), and *tbx6* (f) loci. Scale is number of reads. Each track represents one biological replicate at tailbud stage. **g**) Venn diagram of genes with regions of significant Gon4l-enrichment within gene bodies (blue) or promoters (pink), and genes differentially expressed in MZ*udu*-/- gastrulae (gray). **h**) Correlation of Gon4l enrichment levels across gene bodies (blue dots) and promoters (red dots) with relative transcript levels of genes differentially expressed in MZ*udu* mutants. A positive correlation was detected between increased expression in WT relative to MZ*udu* mutants and Gon4l enrichment in both gene bodies (Spearman correlation p=0.0095) and promoters (p=0.0098). Dotted lines are linear regressions of these correlations.

### Gon4l regulates notochord boundary straightness by limiting *epcam* and *itga3b* expression

We next examined our list of putative direct Gon4l target genes for those with potential roles in tissue boundary formation and/or cell polarity. The *epcam* gene, which encodes Epithelial cell adhesion molecule (EpCAM), stood out because it was not only enriched for Gon4l by DamID (Fig.7a) and upregulated in MZ*udu-/-* gastrulae (Fig.7c), but was also identified in a *Xenopus* overexpression screen for molecules that disrupt tissue boundaries^42^. Furthermore, EpCAM negatively regulates Cadherin-based adhesion^43, 44^ and non-muscle Myosin activity^45^, making it a compelling candidate. We also chose to examine *itga3b,* which encodes Integrinα3b, because as a component of a Laminin receptor^46^, it is an obvious candidate for molecules involved in formation of a tissue boundary at which Laminin is highly enriched (Fig.2f). *itga3b* expression was increased in MZ*udu-/-* gastrulae by both RNA-seq and qRT-PCR (Fig. 7d), and DamID revealed a region within the *itga3b* promoter at which Gon4l was highly enriched (Fig. 7b-b’). We found that overexpression of either *epcam* or *itga3b* by RNA injection into WT embryos recapitulated some aspects of the MZ*udu-/-* gastrulation phenotype, including irregular notochord boundaries (Fig.7e) and reduced ML orientation and elongation of axial mesoderm cells (Fig.7h-m). Conversely, injection of a translation-blocking *itga3b* MO (MO1-itga3b^47^) at a dose phenocopying the fin defects of *itga3b* mutants^47^ (Supplemental Fig.6b) significantly improved boundary straightness in MZ*udu* mutants compared with control injected siblings (Fig.7f). However, injection of an *epcam* MO (MO2-epcam^48^) at a dose phenocopying the loss of otoliths in *epcam* mutants^48^ (Supplemental Fig.6d) did not suppress boundary defects in MZ*udu* mutants (Fig.7g), indicating that excess *itga3b* is largely responsible for irregular notochord boundaries in MZ*udu-/-* gastrulae. To determine whether excess *itga3b* was sufficient to phenocopy the *kny-/-;udu-/-* double mutant phenotype identified in our synthetic screen (Fig.1), we overexpressed *itga3b* in *kny-/-* embryos and found that it exacerbated the short axis phenotype of these PCP mutants at 24hpf (Fig.7n-p), an effect not produced by injection of control *GFP* RNA. Together, these results indicate that negative regulation of *itga3b* and *epcam* expression by Gon4l contributes to proper notochord boundary formation in zebrafish gastrulae. Interestingly, ML cell polarity defects were not suppressed in MZ*udu-/-* embryos injected with *itga3b* (or *epcam)* MO compared to control injected mutant siblings (Supplemental Fig.6e-p), despite the improvement in boundary straightness. This implies that Gon4l has boundary-dependent and independent roles in cell polarization, and that additional gene expression changes likely contribute to this aspect of the mutant phenotype.

**Figure 7:**
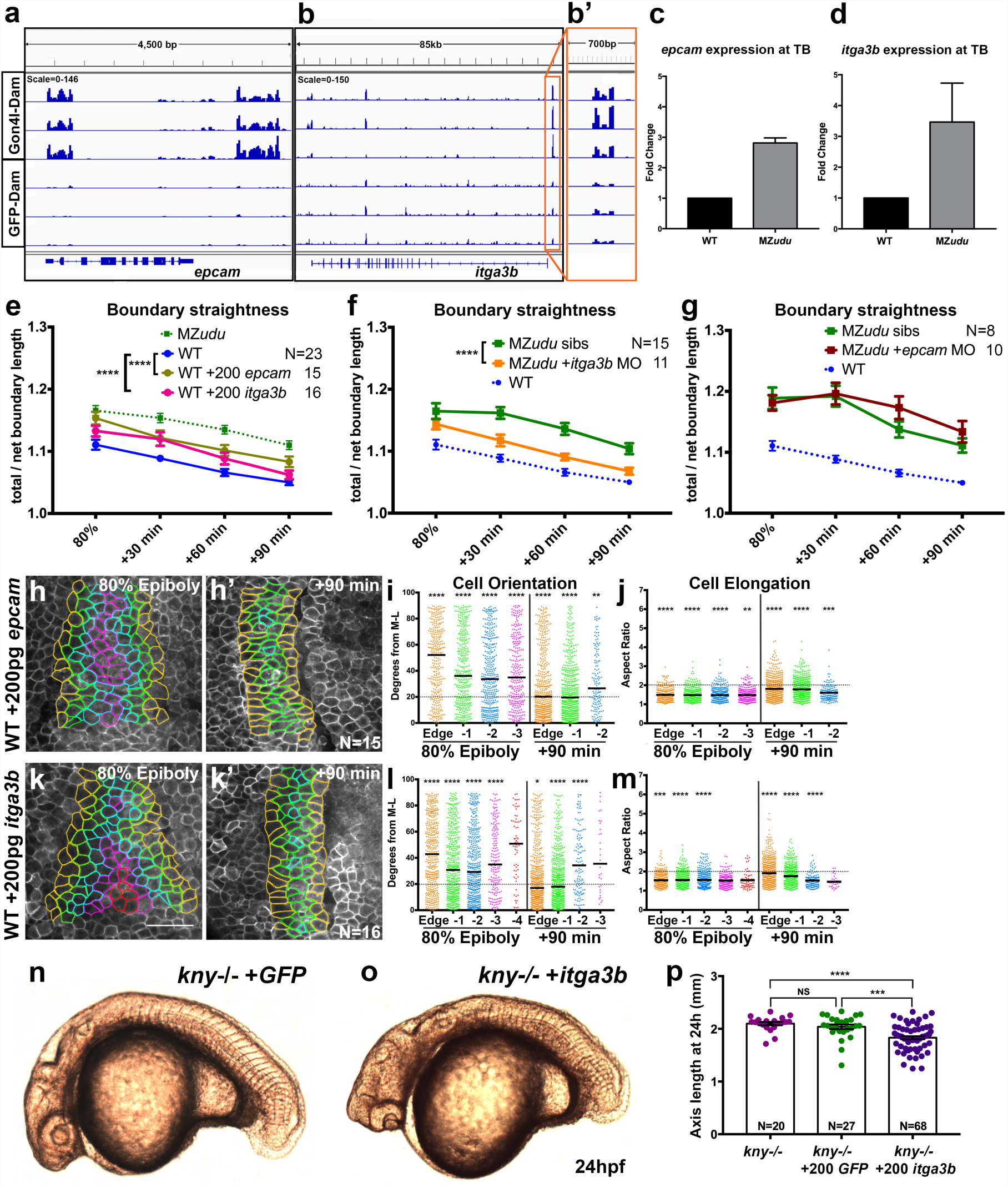
Gon4l regulates notochord boundary straightness by limiting *itga3b* and *epcam* expression. **a-b’)** Genome browser tracks of Gon4l-Dam and GFP-Dam association at the *epcam* (a) and *itga3b* (b) locus. An expanded view of the *itga3b* promoter is shown in b’. **c-d)** Quantitative RT-PCR for *epcam* (c) and *itga3b* (d) in WT and MZ*udu*-/- embryos at tailbud stage. Bars are means with SEM. **e)** Quantification of notochord boundary straightness in WT *epcam* and *itga3b* overexpressing embryos (2-way ANOVA, ^****^p<0.0001). **f-g)** Quantification of notochord boundary straightness in MZ*udu*-/- *epcam* (f) and *itga3b* (g) morphants and sibling controls (2-way ANOVA, ^****^p<0.000l). N indicates the number of embryos analyzed, symbols are means with SEM. h,h’,k,k’) Still images from live time-lapse confocal movies of the axial mesoderm in *epcam* (h) and *itga3b* (k) overexpressing WT gastrulae at the time points indicated. Cell outlines are colored as in Figure 3. **i,l**) Quantification of axial mesoderm cell orientation as in Figure 3. **j,m)** Quantification of axial mesoderm cell elongation as in Figure 3. Asterisks indicate significant differences compared to WT controls. Scale bar is 50μm. **n-o)** Live images of *kny-/-* embryos at 24hpf injected with 200pg *GFP* (n) or 200pg *itga3b* (o) RNA. **p**) Quantification of axis length of injected *kny-/-* embryos at 24hpf. Each dot represents one embryo, bars are means with SEM (T-tests, ^***^p<0.001, ^****^p<0.0001).

### Tissue tension is reduced at the notochord boundary of MZ*udu* embryos

In WT zebrafish and *Xenopus* embryos, the notochord boundary straightens over time (Fig.2e) and accumulates myosin^49^, implying that it is under tension. Mechanical forces within the embryo (like tension) can instruct specific cell behaviors such as polarization, intercalation, and PCP protein localization^50^^-^^54^, and we hypothesized that increased notochord boundary tension may act as an instructive cell polarity cue that is reduced in MZ*udu-/-* gastrulae. We therefore laser-ablated interfaces between axial mesoderm cells in live WT and MZ*udu-/-* gastrulae and recorded recoil of adjacent cell vertices as a measure of tissue tension^55, 56^ (Fig.8a-b). Cell interfaces were classified according to convention described in (^55, 57^) as V junctions (actively shrinking to promote cell intercalation), T junctions (not shrinking) or Edge junctions (falling on and thus comprising the notochord boundary). We found that recoil distances at WT Edge junctions were significantly greater than that of WT V (p=0.0001) or T junctions (p=0.0026), demonstrating that the notochord boundary is under greater tension than the rest of the tissue (Fig.8c-e). Consistent with our hypothesis, recoil distance was significantly smaller in MZ*udu-/-* than in WT gastrulae for all classes of junctions (Fig.8c-e), and this decrease was largest and most significant in Edge junctions, especially at 80% epiboly (Fig.8c). These results point to reduced tension as a possible cause of notochord boundary and cell polarity defects in MZ*udu* mutants. Because excess *epcam* and *itga3b* were sufficient to disrupt notochord boundaries and ML cell polarity (Fig.7), we next tested whether *epcam* and *itga3b* overexpression could likewise affect boundary tension. Injection of WT embryos with *epcam* RNA was sufficient to disrupt tissue tension at the notochord boundary and throughout the axial mesoderm (Fig.8c-e). Curiously, although we found *itga3b* to be largely causative of boundary irregularity in MZ*udu* mutants (Fig.7), its overexpression did not consistently produce a similar reduction in tension (Fig.8c-e). Together these results implicate excess EpCAM and Integrinα3b as key molecular defects underlying reduced boundary tension and reduced boundary straightness, respectively, observed in MZ*udu*-/- gastrulae. We propose a model whereby Gon4l negatively regulates expression of these adhesion molecules to ensure proper formation of the notochord boundary, which together with additional boundary-independent roles of Gon4l cooperates with PCP signaling to promote ML cell polarity underlying C&E gastrulation movements (Fig.8f).

**Figure 8:**
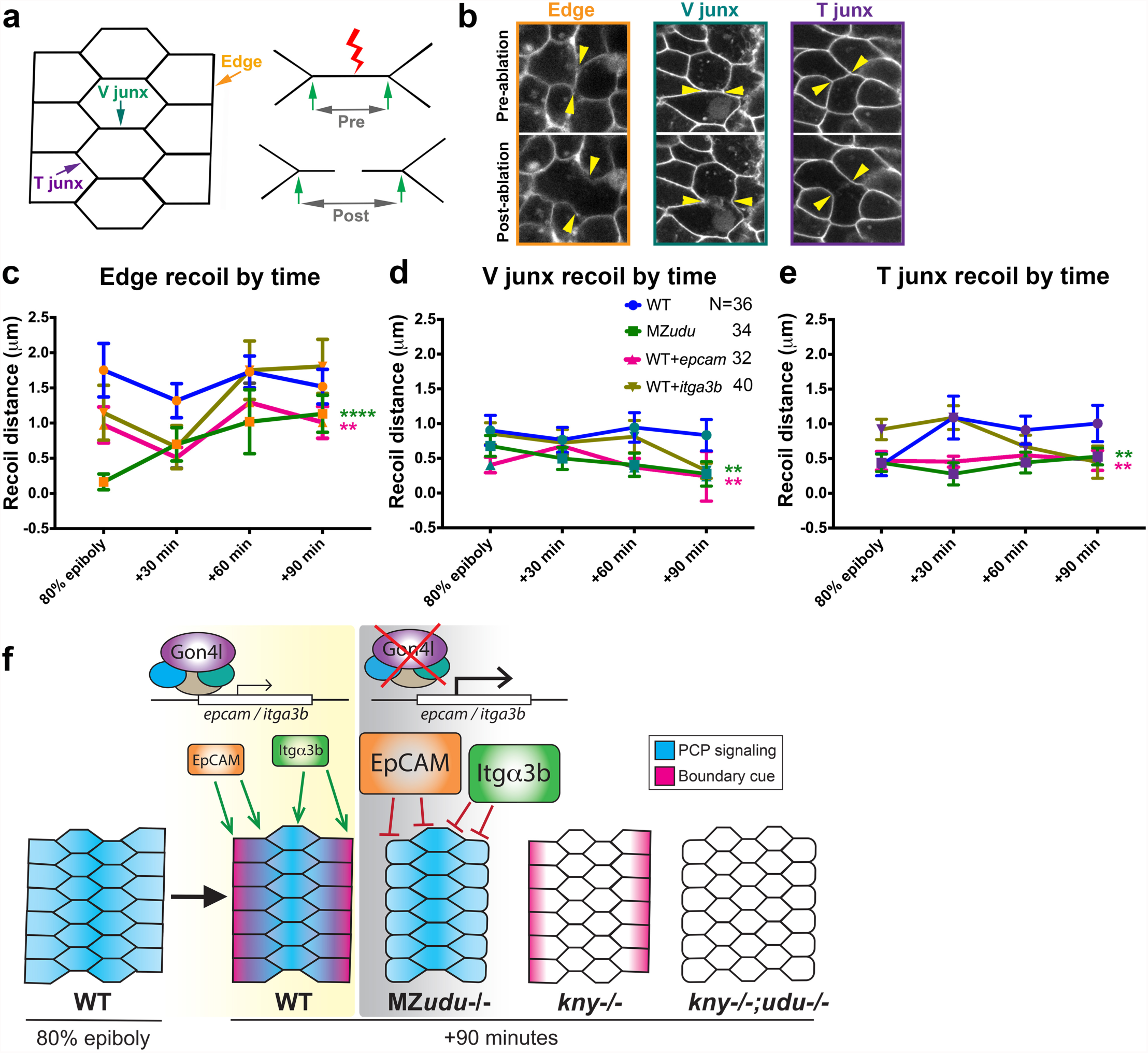
Tension is reduced at the notochord boundaries of MZ*udu* mutant and *epcam* overexpressing WT gastrulae. **a)** Diagram of laser ablation experiments to measure tension at axial mesoderm cell interfaces. **b)** Still images from confocal time-lapse movies of each of the three types of cell interfaces (Edge, V junctions, and T junctions) before and after laser ablation. Arrowheads indicate cell vertices adjacent to the ablated interface. **c-e**) Quantification of cell vertex recoil distance immediately after laser ablation of Edge (c), V junction (d), and T junction (e) interfaces at the time points indicated in WT, MZ*udu -/-,* WT *epcam* overexpressing, and WT *itga3b* overexpressing gastrulae. Symbols means with SEM. Asterisks are colored according to key and indicate significant differences compared to WT controls (2-way ANOVA,^****^ p<0.0001, ^**^ p<0.01). N indicates the number of embryos analyzed. **f**) Graphical model of the roles of Gon4l and PCP signaling in regulating ML cell polarity of axial mesoderm cells.

## DISCUSSION

Significant advances have been made in defining signaling pathways instructing gastrulation cell behaviors that shape the vertebrate body plan, but epigenetic control of these morphogenetic processes remains largely unexplored. Here we have described the conserved chromatin factor Gon4l as a regulator of polarized cell behaviors underlying axis extension during zebrafish gastrulation. We identified a large number of genes regulated by Gon4l, including many genes with known or predicted roles in morphogenesis, and linked misregulation of a subset of them to specific morphogenetic cell behaviors. Because Gon4l does not bind DNA directly, we predict that Gon4l-enriched genomic loci are direct targets of chromatin modifying protein complexes with which Gon4l associates^41^. Only a fraction of the thousands of Gon4l-enriched genes and promoters exhibited corresponding changes in gene expression during gastrulation, which may reflect the following: 1) Gon4l may not alter expression of all loci with which it associates, 2) loci recently occupied by Gon4l may not yet reflect changes in transcript levels, and 3) because DamID provides a “history” of Gon4l association, our experiment revealed both past and currently occupied loci. We also identified loci at which Gon4l was depleted compared to GFP-Dam controls (Supplemental Fig.5), and speculate they represent open chromatin regions with which Gon4l does not associate. Surprisingly, although a larger number of putative Gon4l direct target genes were negatively regulated (i.e. upregulated in MZ*udu* mutants), the highest levels of Gon4l enrichment correlated with positive regulation of gene expression (Fig.6). This demonstrates that Gon4l does not act strictly as a negative regulator of gene expression, a role assigned to it based on *in vitro* evidence and thought to be mediated by its interactions with Histone deacetylases^41^. Our data instead indicate that Gon4l acts as both a positive and negative regulator of gene expression during zebrafish gastrulation, implying context-specific interactions with multiple epigenetic regulatory complexes.

Phenotypes caused by complete *udu* deficiency are conspicuously pleiotropic, but our studies point to remarkably specific roles for *udu* in gastrulation morphogenesis. Loss of *udu* function reduced tissue extension without impairment of other gastrulation movements, including mesendoderm internalization, epiboly, prechordal plate migration, and convergence (see below). Moreover, the dorsal gastrula organizer and all three germ layers were specified (Fig. 1, Supplemental Fig.2), indicating that MZ*udu* mutants do not suffer from a general delay or arrest of development; rather Gon4l regulates a specific subset of gastrulation cell behaviors including ML cell polarity and intercalation in the axial mesoderm. Importantly, the role of Gon4l in these processes is independent of PCP signaling as supported by several lines of evidence, including distinct morphogenetic defects in PCP versus MZ*udu* mutants. Whereas PCP mutants exhibit reduced convergence movements^9, 10, 14^, evidenced by a larger number of cell rows in *kny-/-* axial mesoderm (Fig.4), the axial mesoderm of MZ*udu* mutants contained a normal number of cell rows (Fig.3), indicating no obvious convergence defect. We also observed an intact notochord boundary in PCP mutants, as well as boundary-associated cell polarity (Fig.4). Despite these apparently parallel functions, expression of some PCP genes, including *wnt11, wnt11r, prickle1a, prickle1b, celsr1b, celsr2,* and *fzd2* are regulated by Gon4l as revealed by RNA-seq and DamID-seq experiments (Supplemental Table 1). However, functional redundancies within the PCP network^58, 59^ likely allow for intact PCP signaling in MZ*udu* mutants despite misregulation of some PCP genes. Notably, gastrula morphology and cell polarity defects in MZ*udu* mutants were also distinct from ventralized and dorsalized patterning mutants that exhibit abnormal C&E^13, 15, 60^.

Modulation of adhesion at tissue boundaries has been implicated as a driving force of cell intercalation^49, 61^, and indeed genes annotated as encoding adhesion molecules tended to be expressed at higher levels in MZ*udu*-/- gastrulae (Fig.5). Two of these, *itga3b* and *epcam,* exhibited increased expression in MZ*udu-/-* gastrulae by RNA-seq and qRT-PCR, were identified as putative direct Gon4l targets by DamID (Fig.7), and were each sufficient to disrupt notochord boundary straightness and ML cell orientation and elongation when overexpressed in WT embryos (Fig.7). Furthermore, decreasing levels of Integrinα3b (but not EpCAM) improved boundary straightness in MZ*udu* mutants (Fig.7), while overexpression of *epcam* (but not *itga3b)* was sufficient to reduce tissue tension at the notochord boundary of WT embryos (Fig.8). These observations suggest that excesses of each of these molecules contribute to overlapping and distinct cellular defects. In *Xenopus* gastrulae, EpCAM negatively regulates non-muscle Myosin contractility^45^, and experimental perturbation of Myosin activity disrupts notochord boundary formation^49^. Myosin accumulation also increases tension at tissue boundaries^62, 63^, which biases cell intercalations^52, 54^. This likely explains why excess *epcam* was sufficient to disrupt boundary tension and straightness in WT embryos, although modulation of Integrin-mediated cell-matrix adhesion at the notochord boundary was also necessary to cause MZ*udu* boundary phenotypes. Notably, restoration of boundary straightness in MZ*udu*-/- gastrulae was not sufficient to improve ML cell orientation. This could reflect at least two possibilities: 1) the suppression of boundary phenotypes by the *itga3b* MO was partial and therefor not sufficient to fully restore the boundary-associated cell polarity signal; or 2) the boundary-associated polarity signal was restored, but MZ*udu*-/- axial mesoderm cells were unable to respond to it. Either of these scenarios assumes that additional gene expression changes contribute to reduced ML cell polarity in MZ*udu* mutants, and indeed, several other candidate genes with plausible or known roles in C&E and/or boundary formation were also misregulated in MZ*udu-/-* gastrulae. For example, *epha7,* which encodes an Eph receptor, a class of signaling molecules with well-described roles in tissue boundary formation^64^, was expressed at reduced levels in MZ*udu* mutants. Expression of *daam1,* which encodes a critical link between PCP signaling and Actomyosin contractility^65^, and genes encoding several Myosin light and heavy chain isoforms were also reduced in MZ*udu* mutants (Supplemental Table 1). In addition to excess *itga3b* and *epcam* expression, misregulation of these molecules could also contribute to impaired ML cell polarity in MZ*udu-/-* gastrulae.

In this study of zebrafish Gon4l, we have begun to dissect the logic of epigenetic regulation of gastrulation morphogenesis, and revealed a key role for this chromatin factor in limiting expression of specific genes during gastrulation. This role has important developmental implications, as C&E gastrulation movements are sensitive to both gain and loss of gene function^9, 12^. We propose that by negatively regulating Integrinα3b and EpCAM levels, Gon4l promotes development of the anteroposteriorly aligned notochord boundary and influences ML polarity and intercalation of axial mesoderm cells. In cooperation with PCP signaling, this cue coordinates ML cell polarity with embryonic patterning to drive anteroposterior embryonic axis extension (Fig.8f).

## ACKNOWLEDGEMENTS

We thank Dr. Bas Van Steensel for EcoDam plasmids, Drs. Christine and Bernard Thisse for WISH probes, Dr. Bo Zhang and Dr. Scott Higdon for bioinformatics help, Dr. Matthew Hass for DamID advice, Bisiayo Fashemi for assistance with image analysis, and the Washington University Genome Technology Access Center for library preparation and sequencing services.

## AUTHOR CONRIBUTIONS

M.W., A.S., and L.S.K. designed the study. M.W., A.S., and T.B. performed experiments. C.Y. participated in the initial forward genetic screen. P.G. performed bioinformatic analysis. M.W. and L.S.K. wrote the manuscript. National Institutes of Health grants R01GM55101 and R35GM118179 to L.S.K. and F32GM113396 to M.W., and a W.M. Keck Foundation Fellowship to M.W. in part supported this study.

## METHODS

### Zebrafish strains and embryo staging

Adult zebrafish were raised and maintained according to established methods^66^ in compliance with standards established by the Washington University Animal Care and Use Committee. Embryos were obtained from natural mating and staged according to morphology as described^67^. All studies on WT were carried out in AB^*^ or AB^*^/Tübingen backgrounds. Additional lines used include *kny^m818^, kny^fr6^* ^10^, *udu*^sq1^ ^27^, and *udu^vu66^* (this work). Embryos of these strains generated from heterozygous intercrosses were genotyped by PCR after completion of each experiment. Germline replaced fish were generated by the method described in (^33^). Briefly, donor embryos from *udu^vu66/+^* intercrosses or females with *udu^vu66/vu66^* germline were injected with synthetic RNA encoding *GFP-nos1-3’UTR* ^68^, and WT host embryos were injected with a morpholino oligonucleotide against *dead end1* (MO1-dnd1)^33^ to eliminate host germ cells. Cells were transplanted from the embryonic margin of donor blastulae to the embryonic margin of hosts at sphere stage, and both hosts and donors were cultured in agarose-coated plates. Host embryos were screened for GFP+ germ cells at 36-48 hpf, and the genotype of corresponding donors was determined by phenotype. All putative *udu^vu66/vu66^* germline hosts were raised to adulthood and confirmed by crossing to *udu^vu66/+^* animals prior to use in experiments.

### Synthetic mutant screening

WT male fish were mutagenized by the chemical mutagen N-ethyl-N-nitrosourea (ENU) as described ^22^, then outcrossed to WT females to produce F1 families. F2 families were obtained by crossing F1 fish with fish homozygous for the hypomorphic *knypek* allele *kny^m818^* (rescued by injection with synthetic *kny* WT RNA). F3 embryos obtained from F2 cross were screened by morphology at 12 and 24hpf to identify recessive enhancers of the *kny^m818/m818^* short axis mutant phenotype^10^.

### Positional cloning

We employed the positional cloning approach using a panel of CA simple-sequence length polymorphism markers representing 25 linkage groups^25^ to map the *vu66* mutation to chromosome 16 between the markers of z17403 and z15431. Given phenotypic similarities between *vu66/vu66* and the *udu^sq1/sq1^* mutant phenotype^27^, we sequenced *udu* cDNA from 24hpf *vu66/vu66* mutant embryos revealing a T2261A transversion that is predicted to create Y753STOP nonsense mutation. We designed a dCAPS^69^ marker for the *vu66* mutation and confirmed that no recombination occurred in 810 *vu66/vu66* homozygous embryos.

### Microinjection

One-celled embryos were aligned within agarose troughs generated using custom-made plastic molds and injected with 1-3 pL volumes using pulled glass needles. Synthetic mRNAs for injection were made by *in vitro* transcription from linearized plasmid DNA templates using Invitrogen mMessage mMachine kits. Doses of RNA per embryo were as follows: 100pg *membrane Cherry,* 50pg *membrane GFP,* 25pg *udu-gfp,* 200pg *epcam,* 200pg *itga3b,* 1pg *gfp-dam-myc,* 3pg *udu-dam-myc* for DamID experiments, 20pg *gfp-dam-myc* for MZ*udu* rescue, and 150pg *GFP-nos1-3’UTR* for germline transplantation. To assess Pk-GFP localization, embryos were injected at one cell stage with *membrane Cherry* RNA, then injected with 15pg *Drosophila prickle-GFP* and 20pg *H2B-RFP* RNAs into a single blastomere at 16-cell stage as described^32, 39^. Injections of MOs were carried out as for synthetic RNA. Doses of MOs per embryo were as follows: 3ng MO1-dnd1 ^33^, 4ng MO1-*tri/vangl2* ^38^, 1ng MO2-epcam^48^, 2ng MO1-itga3b^47^.

### Whole-mount in situ hybridization

Antisense riboprobes were transcribed using NEB T7 or T3 RNA polymerase and labeled with digoxygenin (DIG) (Roche). Whole-mount *in situ* hybridization (WISH) was performed as described ^70^.

### Immunofluorescent staining

Embryos were fixed in 4% paraformaldehyde (PFA) in phosphate buffered saline (PBS), rinsed in PBS + 0.1% tween (PBT), digested briefly in 10 mg/mL proteinase K, re-fixed in 4% PFA, rinsed in PBT, and blocked in 2 mg/mL bovine serum albumin + 2% goat serum in PBT. Embryos were then incubated overnight in rabbit anti-Laminin (Sigma L9393) at 1:200, mouse anti-Myc (Cell Signaling 2276) at 1:1000, or rabbit anti-phospho Histone H3 (Upstate 06-570) at 1:500 in blocking solution, rinsed in PBT, and incubated overnight in Alexa Fluor 488 anti-Rabbit IgG, 568 anti-Rabbit, or 568 anti-Mouse (Invitrogen) at 1:1000 in PBT. Embryos were costained with 4’,6-Diamidino-2-Phenylindole, Dihydrochloride (DAPI) and rinsed in PBT prior to mounting in agarose for confocal imaging.

### TUNEL staining

Terminal deoxynucleotidyl transferase (TdT) dUTP Nick-End Labeling (TUNEL) staining to detect apoptosis was carried out according to the instructions for the ApopTag Peroxidase *in situ* apoptosis detection kit (Millipore) with modifications. Briefly, embryos were fixed in 4% PFA, digested with 10 μg/mL proteinase K, refixed with 4%PFA, and post-fixed in chilled ethanol:acetic acid 2:1, rinsing in PBT after each step. Embryos were incubated overnight with TdT and rinsed in stop/wash buffer, then blocked, incubated with anti-DIG antibody, and stained in Roche BM Purple staining solution.

### Microscopy

Live embryos expressing fluorescent proteins or fixed embryos subjected to immunofluorescent staining were mounted in 0.75% low-melt agarose in glass bottomed 35-mm petri dishes for imaging using a modified Olympus IX81 inverted spinning disc confocal microscope equipped with Voltran and Cobolt steady-state lasers and a Hamamatsu ImagEM EM CCD digital camera. For live time-lapse series, 60 μm z-stacks with a 2μm step were collected every three or ten minutes (depending on the experiment) for three hours using a 40x dry objective lens. Embryo temperature was maintained at 28.5°C during imaging using a Live Cell Instrument stage heater. When necessary, embryos were extracted from agarose after imaging for genotyping. For immunostained embryos, 200 μm z-stacks with a 1 or 2μm step were collected using a 10x or 20x dry objective lens, depending on the experiment. Bright field and transmitted light images of live embryos and *in situ* hybridizations were collected using a Nikon AZ100 macroscope.

### Laser ablation tension measurements

Embryos were injected at one cell stage with *mCherry* mRNA and mounted at 80% epiboly for imaging (as described above) on a Zeiss 880 Airyscan 2-photon inverted confocal microscope. An infrared laser tuned to 710 nm was used to ablate fluorescently labeled cell interfaces, immediately followed by a quick time-lapse series of ten images with no interval to record recoil after each ablation event. Images were collected using a 40x water immersion objective lens, and a Zeiss stage heater was used to maintain embryo temperature at 28.5°C. Approximately 8-12 cell interfaces were ablated per embryo, and no fewer than 32 embryos per genotype from no fewer than four independent experiments were analyzed due to high variability of recoil distances. Image series were analyzed using ImageJ to determine the inter-vertex distance of each cell interface prior to and immediately after ablation and used to calculate recoil distance.

### Image analysis

ImageJ was used to visualize and manipulate all microscopy data sets. For immunostained embryos, multiple z-planes were projected together to visualize the entire region of interest. For live embryo analysis, a single z-plane through the length of the axial mesoderm was chosen for each time point. When possible, embryo images were analyzed prior to genotyping. Cell polarity was analyzed in no fewer than six live embryos per genotype due to high variability of these measurements. To measure cell orientation and elongation, the anteroposterior axis in all embryo images was aligned prior to manual outlining of cells. A fit ellipse was used to measure orientation of each cell’s major axis and its aspect ratio. The TissueAnalyzer ImageJ package^71^ was used to automatically segment time-lapse series of axial mesoderm and detect T1 transitions. Boundary straightness was measured by manually tracing the notochord boundary to determine total length, then dividing it by the length of a straight line connecting the ends of the boundary (net length). To assess Pk-GFP localization, isolated cell expressing Pk-GFP were scored according to whether GFP was present in puncta localized to the anterior side of the cell, in puncta localized elsewhere, or in the cytoplasm. Pk-GFP localization was analyzed in multiple cells from each of no fewer than three embryos per genotype.

### Quantitative RT-PCR

Total RNA was isolated from tailbud stage WT and MZ*udu*-/- embryos homogenized in Trizol (Life Technologies), 1μg of which was used to synthesize cDNA using the iScript kit (BioRad) following manufacturer’s protocol. SYBR green (BioRad) qRT-PCR reactions were run in a CFX Connect Real-Time PCR detection system (BioRad). Primers used are as follows:

*epcam*: F- TGAGGACGGGGATTGAGAAC

R- GAGCCTGCCATCCTTGTCAT

*itga3b:* F- CCGGTGTTGGGAGAAGAGAC

R- CTTGAAGAAACCACACGAAGGG

*EF1a:* F- CTGGAGGCCAGCTCAAACAT

R- ATCAAGAAGAGTAGTACCGCTAGCATTAC

*Rpl13a:* F- TCTGGAGGACTGTAAGAGGTATGC

R- AGACGCACAATCTTGAGAGCAG

### RNA sequencing and analysis

RNA for sequencing was isolated from 50 WT or MZ*udu* embryos per sample at tailbud stage according to instructions for the Dynabeads mRNA direct kit (Ambion). Embryos from two clutches per genotype (e.g. WT A & B) were collected, then divided in two (e.g. WT A1, A2, B1, and B2) to yield four independently prepared libraries representing two biological and two technical replicates per genotype. Libraries for were prepared according to instructions for the Epicentre ScriptSeq v2 RNA-seq Library preparation kit (Illumina). Briefly, RNA was enzymatically fragmented prior to cDNA synthesis. cDNA was then 3’ tagged, purified using Agencourt AMPure beads, and PCR amplified, at which time sequencing indexes were added. Indexed libraries were then purified and submitted to the Washington University Genome Technology Access Center (GTAC) for sequencing using an Illumina HiSeq 2500 to obtain single-ended 50bp reads. Raw reads were mapped to the zebrafish GRCz10 reference genome using STAR (2.4.2a)^72^ with default parameters. FeatureCounts (v1.4.6) from the Subread package^73^ was used to quantify the number of uniquely mapped reads (phred score≥10) to gene features based on the Ensembl annotations (v83). Significantly differentially expressed genes were determined by using DESeq2 in the negative binomial distribution model^74^ with a cutoff of adjusted p-value≤0.01 and fold-change≥2.0. Heatmaps were built using the heatmaps2 package in R, and other plots were built using the ggplot2 package in R.

### DamID-seq

*E. coli* DNA adenine methyltransferase (EcoDam) was cloned from the pIND(V5)EcoDam plasmid^21^ (a kind gift from Dr. Bas Van Steensel, Netherlands Cancer Institute) by Gibson assembly into a 3’ Gateway entry vector containing 6 Myc tags and an SV40 polyA signal (p3E-MTpA). The resulting p3E-EcoDam-MTpA vector was then Gateway cloned into PCS2+ downstream of *udu* cDNA or *eGFP* to produce C terminal fusions. The resulting *udu-dam-myc* and *gfp-dam-myc* plasmids were linearized by KpnI digestion and transcribed using the mMessage mMachine SP6 *in vitro* transcription kit (Ambion). WT AB^*^ embryos were injected at one cell stage with 3pg mRNA encoding Gon4l-Dam-Myc or with 1pg encoding GFP-Dam-Myc as controls. The remainder of the protocol was carried out largely as in (^21^) with modifications. Genomic (g)DNA was collected at tailbud stage using a Qiagen DNeasy kit with the addition of RNAse A. gDNA was digested with Dpnl overnight, followed by ligation of DamID adaptors. Un-methylated regions were destroyed by digestion with DpnII, and then methylated regions were amplified using primers complementary to DamID adaptors. Two identical 50μl PCR reactions were performed and pooled for each sample to reduce amplification bias, and three biological replicates were collected per condition. Amplicons were purified using a Qiagen PCR purification kit, then digested with DpnII to remove DamID adaptors. Finally, samples were purified using Agencourt AMPure XP beads and submitted for library preparation and sequencing at the Washington University GTAC using an Illumina HiSeq 2500 to obtain single-ended 50bp reads.

### Analysis of DamID-seq data

Raw reads were aligned to zebrafish genome GRCz10 by using bwa mem (v0.7.12) with default parameters^75^, then sorted and converted into bam format by using SAMtools (v1.2)^76^. The zebrafish genome was divided into continuous 1000bp bins, and FeatureCounts (v1.4.6) from the Subread package^73^ was used to quantify the number of uniquely mapped reads (phred score ≥10) in each bin. Significantly differentially Gon4l-associated bins were determined by using DESeq2 in the negative binomial distribution model^74^ with stringent cutoff: adjusted p-value≤0.01 and fold-change ≥4.0. Promoter regions were defined as 2kb upstream of the transcription start site of a gene based on Ensembl annotations (v83). Wiggle and bigwig files were created from bam files using IGVtools (v2.3.60) (https://software.broadinstitute.org/software/igv/igvtools) and wigToBigWig (v4)^77^, respectively. BigWig tracks were visualized using IGV (v2.3.52)^78^.

### Data Availability

Processed RNA-seq and DamID-seq data are available in Supplemental Table1 of this publication, and raw data have been deposited in the Gene Expression Omnibus (GEO) with the following accession numbers: RNA-seq: GSE96575 https://www.ncbi.nlm.nih.gov/geo/query/acc.cgi?token=ipwzmaoqjpajbqd&acc=GSE96575 DamID-seq: GSE96576 https://www.ncbi.nlm.nih.gov/geo/query/acc.cgi?token=apkloooyvxsjxij&acc=GSE96576

### Statistical analysis

Graphpad Prism 6 and 7 software was used to perform statistical analyses and generate graphs of data collected from embryo images. The statistical tests used varied as appropriate for each experiment and are described in the text and figure legends. All tests used were two-tailed. Differential expression and differential enrichment analysis of RNA and DamID sequencing data were completed as described above. Panther^79^ was used to classify differentially expressed genes and to produce pie charts; DAVID Bioinformatic Resources^80^ was used for functional annotation analysis.

### Subcloning

The full-length *udu* open reading frame was subcloned from cDNA using following primers:

*udu* cacc F1: CACCATGGGATGGAAACGCAAGTCTTC

*udu* TGA R: TCAGTCCTGCTCTTCATCAGTGGC

*udu* R: GTCCTGCTCTTCATCAGTGGCCGAC

*udu* cDNAs with or without a stop codon were cloned into the pENTR/D-TOPO (Thermo Fisher) vector, which were then Gateway cloned into PCS2+ upstream of polyA signal or *eGFP* to produce a C terminal fusion. The full-length *epcam* open reading frame was subcloned from WT cDNA using the following primers:

*epcam* F: GGATCCCATCGATTCGATGAAGGTTTTAGTTGCCTTG

*epcam* R: ACTCGAGAGGCCTTGTTAAGAAATTGTCTCCATCTC

The 5’ portion of the *itga3b* open reading frame was cloned from WT cDNA, and the 3’ portion was cloned from a partial cDNA clone (GE/Dharmacon) using the following primers:

5’*tga3b*F: TGCAGGCGCGCCGGATCCCATCGATTCGATGGCCGGAAAGTCTCTG

5’ *itga3b* R: ATTT GAGTGAGTAT GGAAT G G AGATGTT GAGCG

3’ *itga3b* F: TCAACATCTCCATTCCATACTCACTCAAATACTCAGG

3’ *itga3b* R: GTTCTAGAGGTTTAAACT CGAGAGGCCTTGTCAGAACT CCTCCGT CAG

The resulting amplicons were Gibson cloned^81^ into PJS2 (a derivative of PCS2+) linearized with EcoRI.

